# Inference of enhancer-specific transcription factor interactions from gene expression data using a biophysical model

**DOI:** 10.64898/2026.06.03.729923

**Authors:** Amin Safaeesirat, Hoda Taeb, Eldon Emberly

## Abstract

Transcription factors (TFs) play a central role in gene expression and regulation. In recent years, numerous experimental techniques have generated large-scale datasets, alongside computational methods aimed at inferring the role of TF–TF interactions in gene regulation. However, these approaches typically yield global interaction patterns across datasets, which may not accurately reflect local regulatory interactions at specific enhancers. Here, we model transcription using an Ising-type biophysical framework and introduce approximations based on its mean-field representation to infer TF–TF interactions at the level of individual enhancers from expression data, such as STARR-seq or fluorescent protein measurements. We validate our approach using simulated data and evaluate the effect of the strengths of TF–TF and TF–DNA interactions on inference accuracy. We then apply the model to experimental fluorescence data of gap genes for the eve stripe-2 (eve2) enhancer in the fruit fly embryo. The model successfully infers the established roles of the gap genes and predicts the possibility of cooperative and antagonistic interactions among them, which can be experimentally investigated.

## 1 Introduction

Gene expression begins with transcription, the process by which genetic information encoded in DNA is transcribed into RNA that may then be translated into protein. Precise spatiotemporal control of transcription is essential for proper cellular function, development, and homeostasis. This regulation is primarily mediated by transcription factors (TFs) [1], sequence-specific DNA-binding proteins that modulate gene expression by recruiting or blocking the transcriptional machinery at target loci. The combinatorial action of multiple TFs binding to enhancer sequences along with chromatin structure and other regulatory elements, creates complex regulatory networks that determine when and where genes are expressed. Disruptions to these networks underlie numerous human diseases, including cancer, developmental disorders, and metabolic dysfunction [2].

Advances in sequencing and imaging techniques have led to the ability to measure transcriptional output quantitatively over a population of cells or at the single cell level. Massively parallel reporter assays (MPRAs) [3] enable the simultaneous measurement of expression levels for thousands of regulatory sequences by quantifying the abundance of barcode-associated transcripts in RNA corresponding to each putative enhancer sequence. For example, Self-Transcribing Active Regulatory Region sequencing (STARR-seq) [4, 5] identifies active enhancers by using candidate sequences as their own transcriptional reporters. More broadly, RNA sequencing (RNA-seq) has become a standard approach for genome-wide transcriptome profiling, quantifying expression through read counts mapped to each gene [6]. Fluorescence-based methods, including fluorescent proteins as reporters and quantitative polymerase chain reaction (qPCR) with fluorescent probes, provide measurements of gene expression levels in individual cells or populations [7]. These diverse experimental techniques generate rich datasets capturing regulatory activity across different genomic contexts, sequence variants, and cellular conditions. However, extracting mechanistic insights about transcriptional regulation from these large-scale measurements requires appropriate analytical and modeling frameworks.

Machine learning and deep learning approaches are widely used to analyze transcriptional data. These methods have been applied to tasks such as predicting enhancer activity and gene expression levels directly from DNA sequence [8–12], identifying TF binding motifs [13, 14], and inferring gene regulatory networks [15–17]. By leveraging large-scale datasets and flexible model architectures, they can capture complex, nonlinear dependencies in gene regulation and often achieve strong predictive performance across diverse genomic contexts [18]. A key advantage of such data-driven approaches is their ability to integrate heterogeneous data types and scale to genome-wide analyses. However, they often lack immediate interpretability and are difficult to translate into mechanistic insight. Furthermore, these models are limited in their ability to identify short, low information motifs many TFs possess, and to disentangle the contributions of redundant TFs that collaborate within gene regulatory networks. Effort has therefore been devoted to developing interpretability and attribution methods to better understand learned representations and regulatory logic [15, 19–24].

Biophysical and mechanistic modeling approaches aim to describe transcription in terms of explicit molecular interactions, such as TF binding, chromatin accessibility, and transcriptional kinetics [25–34]. These models are inherently more interpretable and can provide insight into regulatory mechanisms. Nevertheless, gene expression is a highly nonlinear and context-dependent process, and faithfully capturing its complexity can lead to models with many parameters and strong assumptions. As a result, biophysical models often require simplifying approximations that may limit their applicability or predictive power in complex regulatory settings. To address this issue, in our previous work (Ref. [25]), we employed an Ising-type biophysical model of transcription to identify TF binding sites and infer the regulatory roles of TFs at those sites for a given enhancer using STARR-seq data. However, that framework did not capture cooperative or antagonistic interactions between TFs in transcriptional regulation.

Here, we show that when TF-specific information, such as binding-site positions, TF–DNA binding energies, and TF concentrations, is available in addition to transcriptional readouts for a given enhancer, it becomes possible to infer pairwise interactions between TFs. To this end, we build on the Ising-type biophysical model used in our previous work and develop a series of inference methods by introducing approximations to the original formulation. We evaluate the performance of these models using simulated STARR-seq data. We then demonstrate that it is possible to infer the underlying regulatory network by applying one of these models to expression data, such as fluorescent protein measurements, collected under different cellular conditions. Finally, we apply our approach to data for eve2 enhancer [20] as a case study to infer interactions among the gap genes.

## 2 Materials and methods

### 2.1 Fully interacting model for transcription

Our biophysical model of transcription is illustrated in Fig. 1. We consider a DNA enhancer fragment *ν*, containing *N*_*ν*_ − 1 binding sites for a set of known TFs. Each site is assigned a binary variable *σ*_*i*_, indicating whether it is bound (*σ*_*i*_ = 1) or unbound (*σ*_*i*_ = 0) by its corresponding TF. In addition to these binding sites, there is an additional site (p) representing the promoter, where RNA polymerase II (Pol-II) can bind and initiate transcription. Thus, the enhancer fragment comprises a total of *N*_*ν*_ sites.

**Figure 1:**
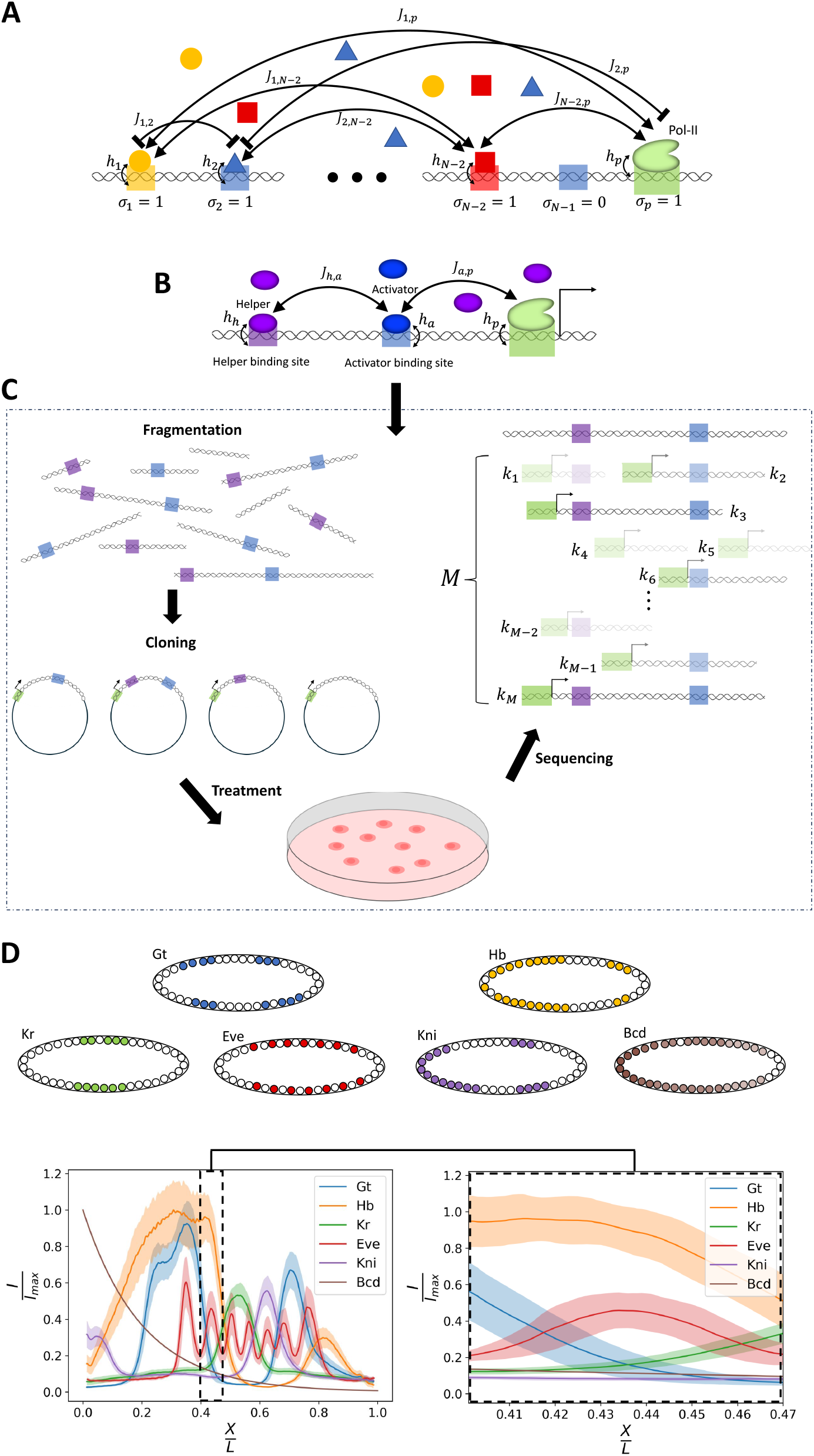
A) Schematic of an enhancer containing different TF-specific binding sites and a promoter, illustrating interaction energies between bound TFs and the promoter. Different TFs can have different concentrations depending on the cellular condition. B) A helper–activator system, shown as an example of a simple enhancer architecture. C) STARR-seq assay applied to the helper–activator system, showing how enhancer activity is experimentally measured. D) Schematic overview of the eve dataset used in this work. The bottom panels show the measured fluorescence intensity of different proteins across the embryo, normalized by the maximum average observed intensity, *I*_max_. In these plots, the solid line represents the mean normalized intensity across the selected embryos, while the shaded region indicates the standard deviation. For further details, see Sec. 2.3.2.

TFs can bind to their specific sites and interact with one another as well as with Pol-II to regulate transcription, as shown in Fig. 1A. The binding of a TF or Pol-II to its site is characterized by an effective binding energy, *h*_*i*_, and a chemical potential 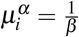 ln 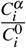, where the cellular condition *α* can affect 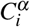 the concentration of TF *i*, relative to a reference concentration, 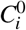. Pairwise interactions between TFs are described by interaction energies *J*_*i j*_, which are assumed to be symmetric (*J*_*i j*_ = *J*_*ji*_) and exclude self-interactions (*J*_*ii*_ = 0). Furthermore, the interaction energies are assumed to be independent of the condition. Given these definitions, the Hamiltonian describing the enhancer can be written as

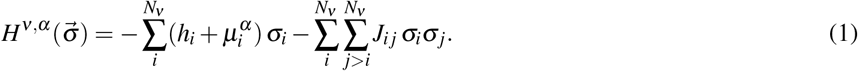

Given the above Hamiltonian and assuming the system is in thermal equilibrium, the probability that the site *i* is occupied is

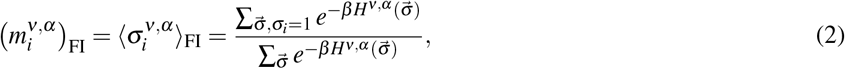

where we take 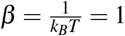. Here, the subscript “FI” represents “the fully interacting model”, in which all pairwise interactions between TFs and Pol-II are included. The above sum requires summing over all possible states of the system, which grows exponentially with the number of TF binding sites in the enhancer.

### 2.2 Mean-field and linear approximations

In the prior section, the equilibrium occupancy of Pol-II required summing over all states, and here we provide several simplifying approximations that make the calculation tractable for systems where enumeration would be infeasible.

Using the fully interacting model (Eq. 2), the binding probability of Pol-II and TFs to their respective sites in fragment *ν* can be described at the mean-field (MF) level by

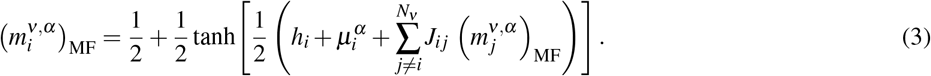

Making the further assumption that the total effective field acting on each site is small, the hyperbolic tangent in Eq. 3 can be linearized using the first-order Taylor expansion (tanh *x* ≈ *x*). This approximation yields

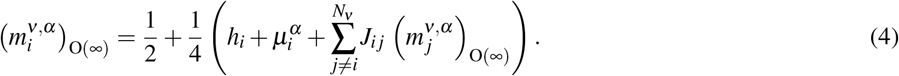

In matrix notation, the solution to Eq. 4 can be written as

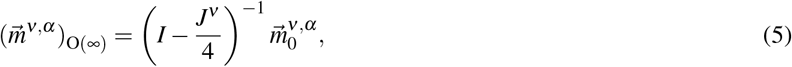

where 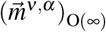 is the vector of site occupancies, *I* is the identity matrix, and *J*^*ν*^ is an *N*_*ν*_ × *N*_*ν*_ symmetric matrix of interaction energies. The vector 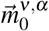 is defined as

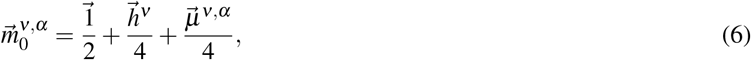

With 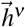 and 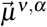 denoting the vectors of binding energies and chemical potentials, respectively. 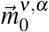 represents the average occupancy of different sites in the absence of the pairwise interactions. Thus, given the system’s parameters, by computing an inverse matrix, one can then find an approximate estimate of the site average occupancies.

The inverse matrix in Eq. 5 can be expanded using the Neumann series,

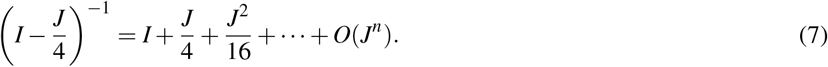

Retaining all terms in this expansion corresponds to the infinite-order model, indicated by the superscript O(∞) in Eq. 5. Truncating the Neumann series at different orders yields a hierarchy of approximate models. Keeping only the first two terms leads to the first-order model,

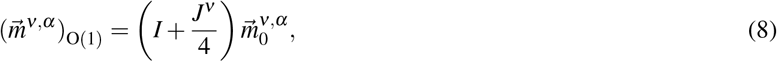

which provides linear approximation for calculating Pol-II occupancy given the system parameters (see Eq. 15).

Binding of Pol-II to its promoter is a prerequisite for transcription. The probability of Pol-II binding can vary across genomic fragments due to differences in TF binding sites, or for a single enhancer, due to cell-to-cell variations in the concentrations of the TFs that interact with it. We assume that the average number of mRNAs (*λ*) produced from an enhancer fragment, *ν*, is proportional to the Pol-II binding probability. So

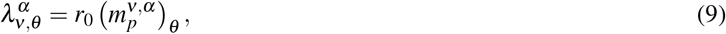

where *r*_0_ is a proportionality constant that captures experimental factors such as measurement duration, sequencing depth, quality of fluorophore, etc., and *θ* represents the model.

As we will see below, our main goal will be to fit the above approximate models to data to find the interaction matrix, *J*. Fitting the mean-field model (Eq. 3) requires optimizing a non-linear system of equations, whereas the last two approximations (Eq. 7 and Eq. 8) lead to simpler computations. Eq. 5 is particularly useful for estimating *J* when the site biding energies, *h*_*i*_, are unknown, but the differences in chemical potentials between two cellular conditions are known.

### 2.3 Transcription data

In this work, we analyze two types of datasets. The first is a simulated dataset comprising mRNA counts from DNA fragments containing different combinations of TF binding sites, assumed to be measured under identical cellular conditions (e.g., STARR-seq, Fig. 1C). The second is a dataset consisting of fluorescence intensity measurements of the expression levels of gap genes at different positions along the Drosophila embryo [20], where the same DNA sequences are exposed to varying cellular environments and TF concentrations (see Fig. 1D). However, our method is not limited to these specific examples and can be broadly applied to similar datasets and experimental contexts.

#### 2.3.1 Self-transcribing active regulatory region sequencing (STARR-seq)

A schematic of the STARR-seq assay [4] is presented in Fig. 1C. In this assay, DNA fragments, either genome-wide or pre-selected by an enrichment method, are cloned into plasmids downstream of a known promoter. If a fragment acts as an enhancer, it drives its own transcription, producing mRNA that contains the fragment’s sequence. This enables the identification of enhancer fragments by sequencing. The plasmids are introduced into cells via electroporation and subjected to different cellular conditions, such as the presence or absence of specific hormones. Finally, sequencing quantifies the mRNA count corresponding to each fragment, providing a quantitative readout of its transcriptional activity. For details on how we simulate STARR-seq data in this work, see Sec. 2.4.

#### 2.3.2 Early embryonic gap gene expression data in Drosophila

We used the wild-type gap gene data provided in [20], where the fluorescence intensity of antibody-labeled gap gene proteins including Hunchback (Hb), Krüppel (Kr), Giant (Gt), and Knirps (Kni), and the even-skipped (eve) protein was measured over time and across spatial positions in the embryo, as illustrated in Fig. 1D. We selected embryos with available fluorescence data between 32 and 36 minutes for the gap genes (28 embryos) and between 40 and 44 minutes for eve (17 embryos), and averaged the signal intensity at different positions for all these embryos. The time difference between the gap gene and eve measurements reflects the approximately 8-minute response delay between changes in the gap gene concentrations and the resulting eve expression [20]. We then focused on data corresponding to the location of the eve2 stripe, defined as the region from *x*/*L* = 0.40 to *x*/*L* = 0.47, where *L* is the length of the embryo (Fig. 1D, zoomed plot), which includes 70 data points. For the fluorescence intensity of the maternal morphogen Bicoid, we assumed an exponential decay across the embryo, as direct measurements were not available in the dataset. The concentration of Bicoid is modeled as

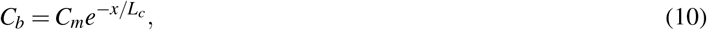

where *x* denotes the position along the anterior-posterior axis of the embryo, *C*_*m*_ is the maximum concentration, and *L*_*c*_ is the characteristic decay length. Based on experimental observations, we take *L*_*c*_ = 0.2*L* = 100 *µ*m [35]. The total length of the embryo is approximately 500 *µ*m.

### 2.4 Simulating STARR-seq data

To simulate STARR-seq data for an enhancer, we assign values to all the TF chemical potentials, binding energies, and pairwise interaction energies, either randomly or according to the regulatory function of interest. Together, these choices result in a system in which the average occupancy of the Pol-II site in Eq. 2, corresponding to the Pol-II binding probability, can be determined. After assigning the interaction energies, we tile the enhancer by generating a set of random overlapping fragments. Each fragment contains a random number of consecutive binding sites drawn from the original enhancer. Fragment length is not explicitly considered in the simulations; only the set of sites present matters. The number of sites in fragment *ν*, denoted by *N*_*ν*_, ranges from 1 (containing only the promoter) to the total number of sites in the enhancer, *N* (all sites present). We assume that the Pol-II site is always present in all fragments, consistent with the STARR-seq assay (Fig. 1C).

Given the binding sites present in a fragment and their associated energies, the probability of Pol-II binding is computed using Eq. 2. For each fragment, the number of produced mRNAs is then drawn from a Poisson distribution with mean 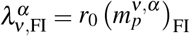 . The resulting mRNA counts across all fragments constitute the simulated dataset.

### 2.5 Fitting models to determine interaction energies, *J*

#### 2.5.1 Likelihood-based inference of model parameters from Pol-II binding probability

For the data that is measured by sequencing, where the output is a count, we assume that the number of observed mRNA molecules for a given fragment follows a Poisson distribution,

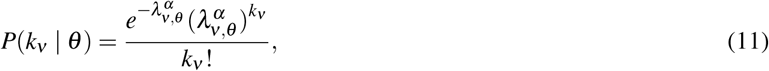

where *k*_*ν*_ shows the number of transcribed mRNA molecules from fragment *ν*. Assuming statistical independence between fragments, the likelihood of the dataset 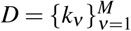 with M fragments given the model is

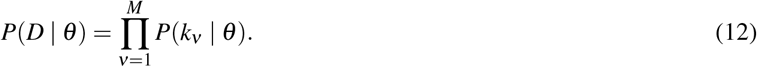

So the log-likelihood takes the form

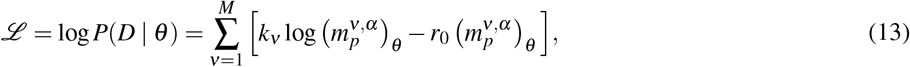

where terms independent of the model have been omitted. Here, the optimal model parameters are obtained by maximizing the log-likelihood using numerical optimization methods, including Monte Carlo sampling and gradient-based optimizers implemented in `SciPy` [36].

For the data where the output is a continuous variable (such as fluorescence intensity), we assume that the data has Gaussian fluctuations around the expected value, 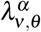. In this case, rather than using Eq. 13, we employ a mean squared error loss function,

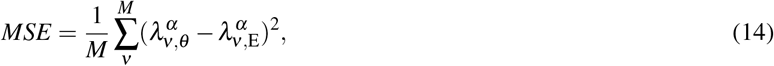

where 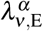 is the average observed expression level in the experiment. Here, *ν* represents an enhancer within an organism, and *α* denotes a specific cellular context that can vary from one cell to another within the organism.

#### 2.5.2 Fitting models to data

We take four different approaches by fitting the biophysical models described above to the transcriptional data in order to estimate the interaction energy matrix, *J*. Positive *J*_*i j*_ indicates that factors *i* and *j* tend to cooperate to activate transcription when bound simultaneously to their sites, whereas a negative entry indicates a repressive interaction or an antagonistic role in transcription.

The first approach is to directly use Eq. 2 to find the set of interaction energies that reproduce the observed Pol-II binding probability, 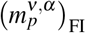, from the data. This can be achieved through Monte Carlo simulations or other minimization techniques, where the optimum set, 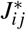, which gives the maximum likelihood of data (Eq. 12) or the minimum error (Eq. 14), can be found. The second approach is based on the mean-field approximation of the model (Sec. 2.2). Eq. 3 can be written separately for each of the *N*_*ν*_ sites within a given fragment, resulting in a system of *N*_*ν*_ coupled nonlinear equations, that given an interaction matrix, *J*_*i j*_, can be solved for unknown 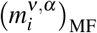. Starting with an initial guess for *J*_*i j*_, one can iteratively update *J*_*i j*_ by minimizing the loss functions.

The third approach employs the linearized version of the mean-field model (Eq. 5), to which a similar iterative procedure as in the full mean-field method can be applied. However, rather than solving a system of *N*_*ν*_ nonlinear equations at each iteration, this approach requires only the computation of a matrix inverse.

The fourth approach is to use the first order model (Eq. 8). For a given fragment, *ν*, the average occupancy of Pol-II can be written as

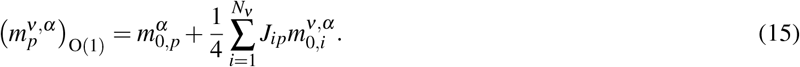

This model does not directly provide the interaction energies, *J*_*i j*_, instead it can estimate the effective interactions with Pol-II, which represent the contribution of a TF to transcription [25]. We perform a Poisson regression fit using this model (see S.M. 1.2).

### 2.6 Fluorescent intensity and chemical potential

At a given cellular position, *x*, in an organism, we assume that the measured fluorescence intensity of a protein is proportional to its local concentration. Under this assumption, the chemical potential of protein *i*, can be written as

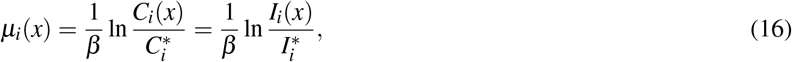

where *C*_*i*_(*x*) and *I*_*i*_(*x*) show the concentration and fluorescence intensity of TF *i* at position *x*, respectively, and 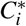 and 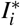 denote the corresponding reference concentration and intensity for TF *i*.

Choosing the reference concentrations is arbitrary. We set them such that they yield the Pol-II binding probability closest to 0.5 in the data, since our approximation is most accurate around this value for the O(∞) model (see Eq. 4). Furthermore, we set *β* = (*k*_*B*_*T*)^−1^ = 1 in the simulations (Sec. 3.3), while treating it as a free (rescaling) parameter in the analysis of the eve2 enhancer data.

## 3 Results

### 3.1 Inference of TF interaction energies from simulated STARR-seq data for simple enhancer architectures

We begin our analysis by first inferring the interactions between a set of known TFs that regulate a set of enhancer sequences for which the transcriptional output has been measured. For a given enhancer sequence, *ν*, the binding site locations and binding energies, *h*_*i*_, and chemical potentials, *µ*_*i*_, are known. Assuming that the measured output is proportional to the equilibrium occupancy of the Pol-II binding site (see Fig. 1C), we have derived several models of decreasing computational complexity, from complete enumeration of all the interacting bound states, to linearized mean-field model, O(∞), and the linear model, O(1) (see Methods) that can learn the interaction energies, *J*_*i j*_, between all the TFs and the TFs and Pol-II. For a sequence with *N*_*ν*_ binding sites, there are 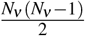 interaction energies to determine, and thus the data requires at least this many independent measurements to be fully determined.

We consider several simple enhancer architectures (see [25, 37]) and simulated data for them akin to that which might be generated by a STARR-seq assay (see Fig. 1C). The first enhancer contains two TF binding sites: one for a helper and one for an activator (see Fig. 1B and Fig. 2A). The activator increases the binding probability of Pol-II, whereas the helper facilitates activator binding. There is no direct interaction between the helper and Pol-II; instead, the helper influences Pol-II binding indirectly by promoting activator occupancy. The second enhancer (Fig. 2B) consists of a repressor, which decreases the binding affinity of Pol-II, and an inhibitor, which prevents binding of the repressor. In this system, the inhibitor indirectly promotes Pol-II binding by suppressing its repressor. The third enhancer (Fig. 2C) contains two repressors. Each repressor can directly decrease Pol-II binding affinity. In contrast to the previous systems, these two repressors also exhibit an activating interaction with each other, which increases the binding affinity of both sites.

**Figure 2:**
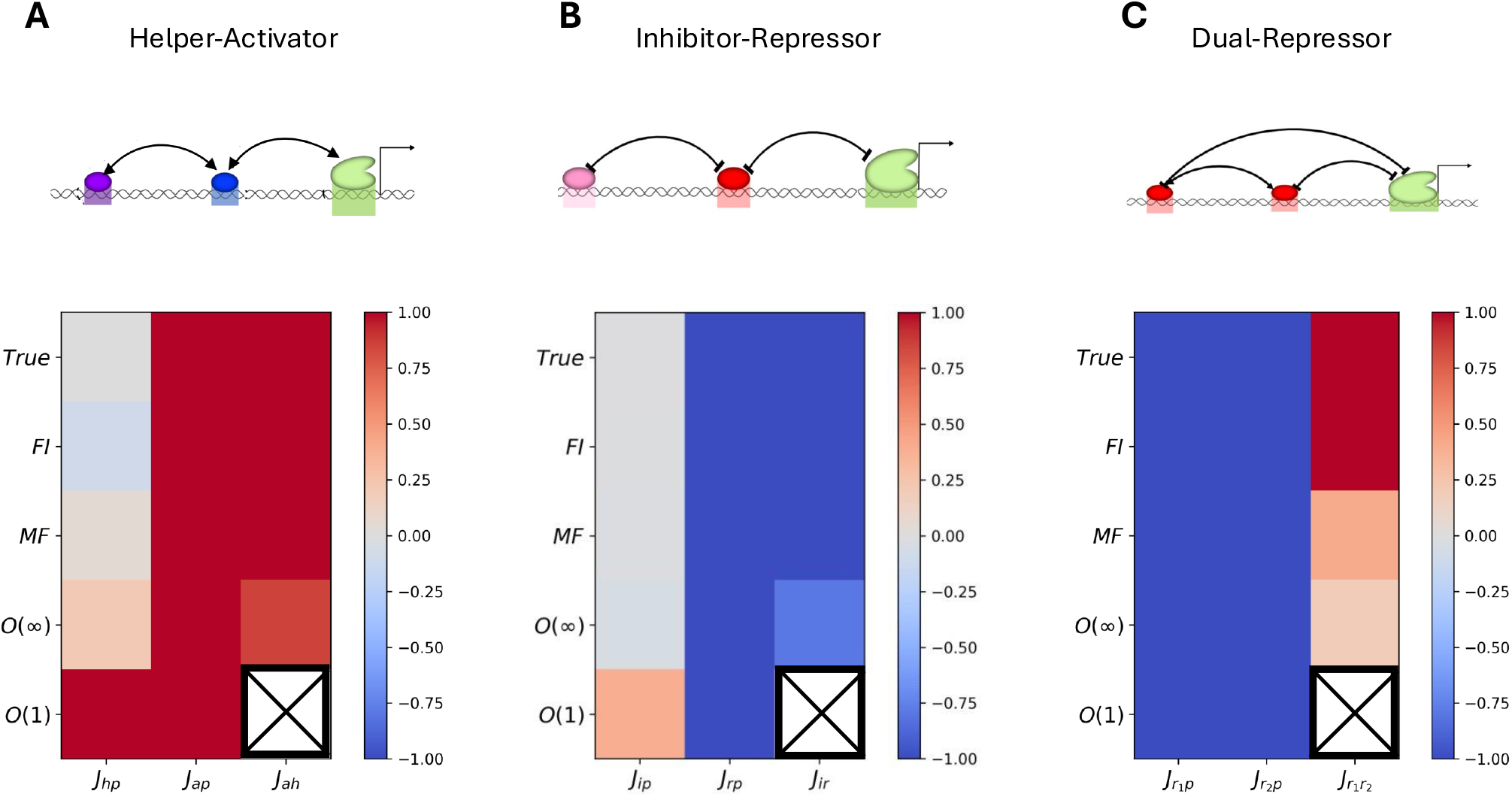
Ground-truth and inferred interaction energies across different models for A) helper–activator, B) inhibitor–repressor, and C) dual-repressor systems. In the first-order model (O(1)), pairwise interaction energies between TFs are not inferred and are therefore indicated by a “×”.

We simulated STARR-seq data for all three systems using *M* = 200 fragments (see Sec. 2.4). The true energy values used in the simulations are listed in S.M. 1.1. Using the four different inference models, we estimated the underlying interaction energies, *J*_*i j*_. The inferred energies for each enhancer and model, together with the ground-truth energies used in generating the data, are shown in Fig. 2. As can be seen, all methods except the first-order linear model (O(1)) correctly recovered the sign of the interaction energies, which reflects the role of each interaction in transcription. However, the magnitude of the inferred energies varies in accuracy across methods. Overall, the FI and MF models perform better than the O(∞) model for these small systems, which is expected as the O(∞) model is a linearized version of the MF model.

Unsurprisingly, the first order linear model, O(1) cannot learn the pairwise interactions. Instead, it infers the *effective* interaction of each factor with Pol-II[25]. For example, in the helper–activator system, even though the helper does not directly interact with Pol-II, the model infers a positive (activating) effective interaction energy, correctly capturing the helper’s indirect activating role. Similarly, in the inhibitor–repressor system, it assigns a positive effective interaction to the inhibitor, reflecting its indirect activation of Pol-II by suppressing the repressor. In all three enhancer architectures, the linear model captures the functional role of each factor, even though it cannot estimate explicit TF–TF interaction energies.

### 3.2 Inference of TF interaction energies from simulated STARR-seq data for complex enhancers

For enhancers containing significantly more binding sites for TFs, as is the case for many eukaryotic enhancers, inferring interaction energies, *J*_*i j*_, from STARR-seq data becomes substantially more challenging. The fully interacting (FI) model is computationally prohibitive as it requires enumeration of a large state space (see Methods). The first-order model (O(1)), on the other hand, is inadequate because it only captures effective interactions with Pol-II and cannot resolve TF–TF interaction energies. Consequently, the most practical approach for enhancers with many TF binding sites is the mean-field (ML) and the all-order (O(∞)) model. In this section, we apply these two models in addition to the FI model to simulated data from larger enhancers to assess their performance in inferring TF–TF interactions.

We constructed enhancers containing seven binding sites, including the promoter site. The interaction energies were drawn randomly from a normal distribution with mean zero and standard deviation *σ* . STARR-seq data were then simulated following the procedure described in Sec. 2.4 for *M* = 1000 fragments (see S.M. 1.1). All the three models were subsequently fit to the simulated data to infer the interaction matrix *J*_*i j*_.

Fig. 3A shows an example heatmap of the true interaction energies used in the simulations alongside the inferred values of *J*_*i j*_ across different models, where *σ* = 1. In this example, the Pearson correlation coefficient between the true and inferred values is 0.98, 0.94, and 0.84 for the FI, MF, and O(∞) models, respectively, indicating strong agreement.

**Figure 3:**
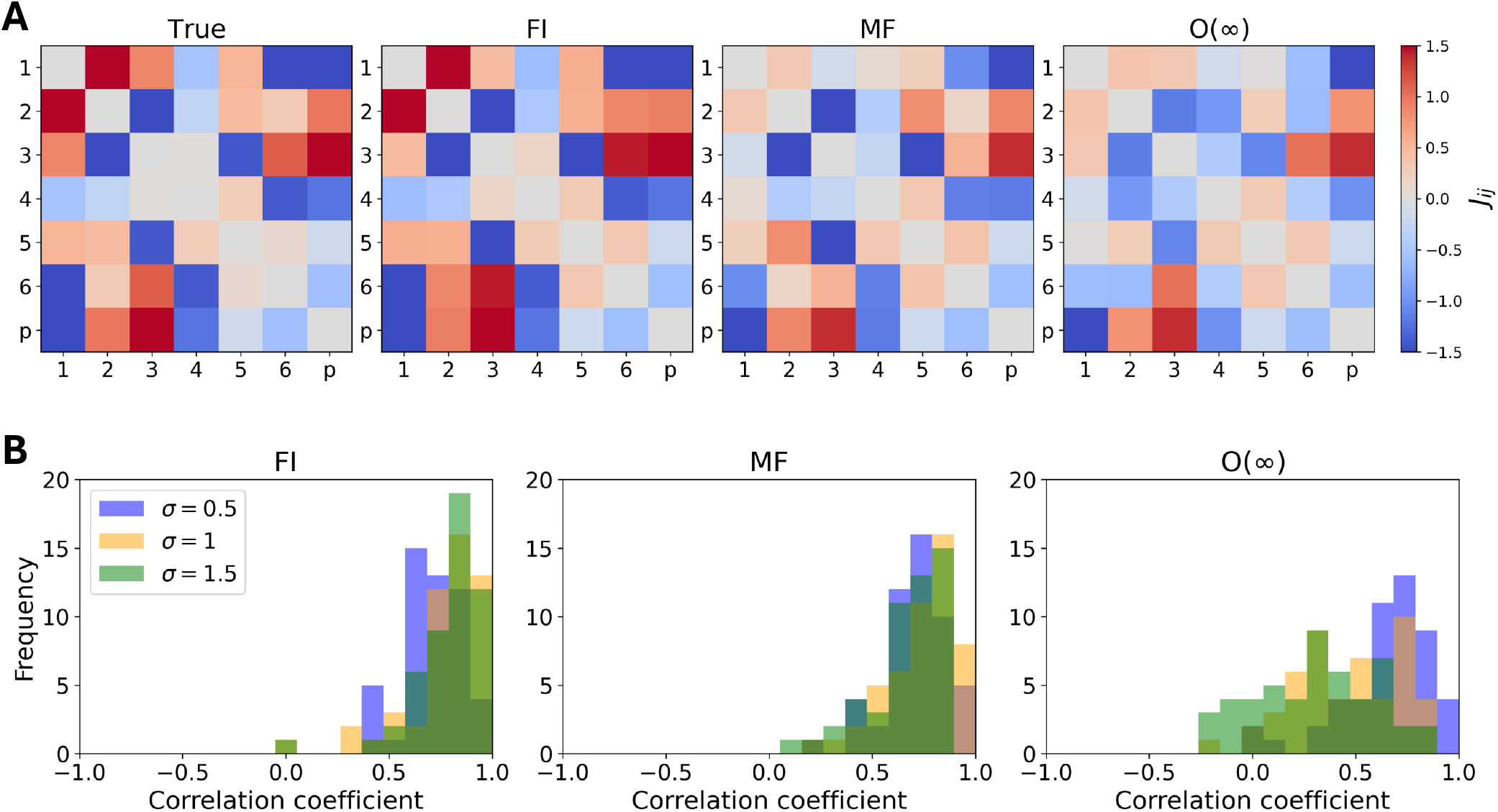
Inference of interaction energies for complex enhancers (six TF binding sites (1-6) and Pol-II(p)) using *M* = 1000 random fragments for each enhancer. A) Heatmaps of the true interaction energy matrix for a particular enhancer alongside the inferred interaction energies using the FI, MF, and O(∞) models, where *σ* = 1. The Pearson correlation between the inferred and true energies is 0.98, 0.94, and 0.84 for the FI, MF, and O(∞) models, respectively. B) Distributions of Pearson correlation coefficients between inferred and true interaction energies across 50 random systems with seven binding sites, for *σ* = 0.5, 1, 1.5.

To assess significance of the fits, this procedure, including the generation of random interaction energies, was repeated 50 times for each value of *σ* . Fig. 3B shows the distribution of correlation coefficients between the true and inferred interaction matrices. As can be seen, the correlation is not consistently high and can vary from weak to strong depending on the underlying random interactions. The performance of the FI and MF models does not change considerably with increasing *σ* . However, for the O(∞) model, the average correlation decreases as *σ* increases, reflecting the reliance of the infinite-order model on the assumption of weak interactions within the system.

To compare these distributions with a random background distribution, we generated 10000 random symmetric matrices with the same *σ* values as the true interaction matrices shown in Fig. 3B and Fig. S1A-B, and calculated the correlation between the true interaction matrices and the randomly generated ones ((Fig. S1C)). As expected, the means of the background distributions are around zero. In contrast, the means of the inferred distributions remain above zero, although they approach zero as *σ* increases in the O(∞) model. This means that the high correlations between the true and inferred interactions are not accidental.

In the above simulations, we considered data measured for a single enhancer in a single condition where it was segmented into fragments. Here we consider the measurement in a single condition (i.e. fixed *µ*_*i*_) of many different enhancers regulated by a known set of TFs. Therefore, TFs interact with the same interaction matrix *J* across different enhancers. The binding energies (*h*_*i*_) vary from one enhancer to another, resulting in strong and weak sites for a given factor across enhancers. Given a sufficient variety of enhancer sequences, and their corresponding fragments, the interaction energies can be inferred using the various approximations as we now show.

We considered 50 enhancers, each randomly tiled 20 times, giving a total of *M* = 1000 data points, matching the sample size of the previous case. Fig. 4A shows the inferred *J* and chemical potentials (the lowest row of the heatmaps) across different models for an example with random *J* and chemical potentials generated with *σ* = 1. In this example, the Pearson correlation coefficient between the true and inferred values is 0.96, 0.97, and 0.71 for the FI, MF, and O(∞) models, respectively. As can be seen, the performance of all three models has improved on average for this data type compared to the previous case (single enhancer with *M* = 1000 fragments). A similar trend is also observed: increasing *σ* worsens the performance of the O(∞) model, however slightly improving the performance of the FI and MF models. For background distributions and heatmap examples for *σ* = 0.5 and *σ* = 1.5, see Fig. S2.

**Figure 4:**
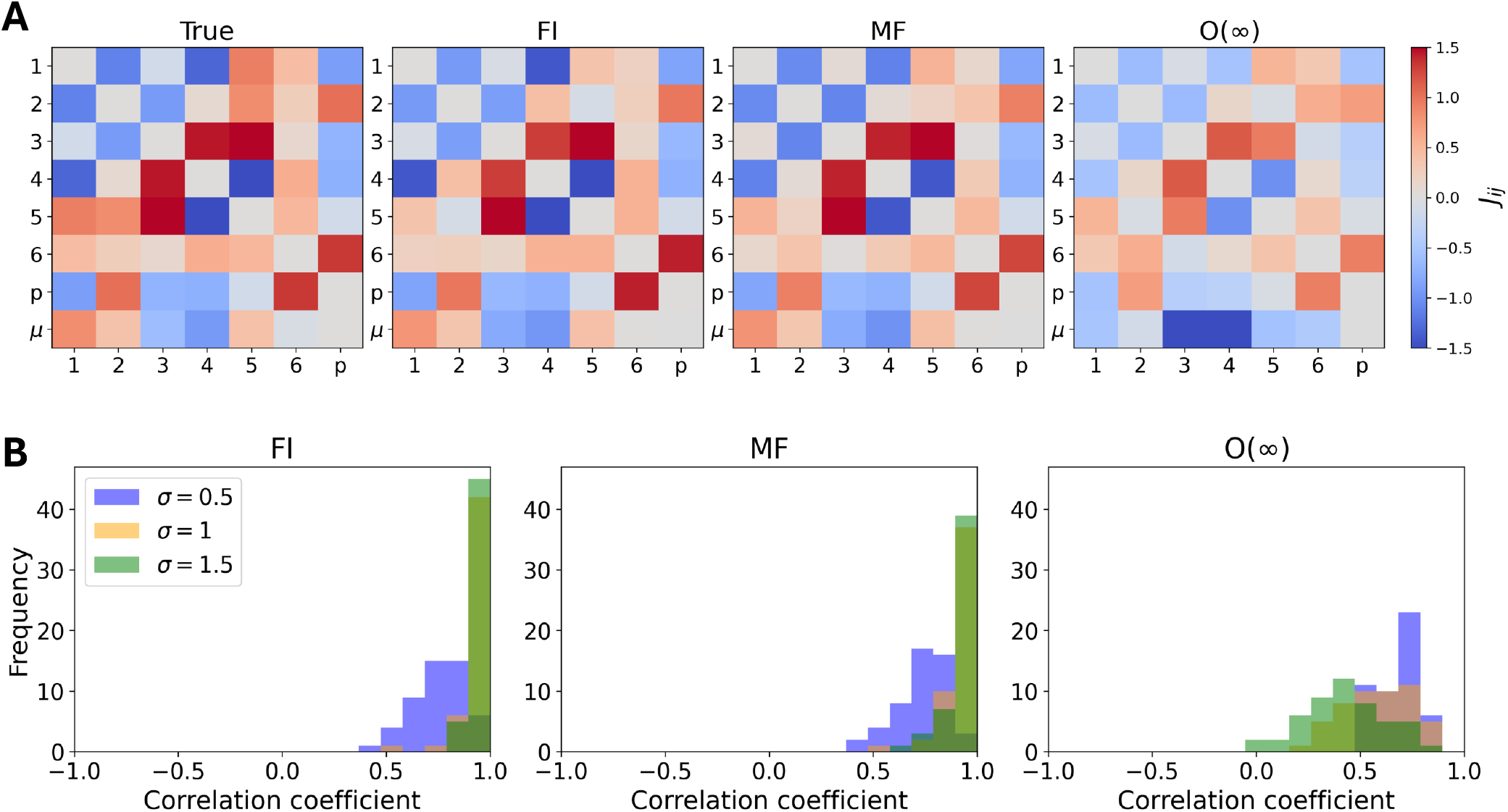
Inference of interaction energies from systems with multiple enhancers (n=50), where each enhancer has varying-strength binding sites for the same TFs, and is randomly tiled with 20 fragments. A) Heatmaps of the true interaction energy matrix alongside the inferred interaction energies and chemical potentials using the FI, MF, and O(∞) models, where *σ* = 1. The Pearson correlation between the inferred and true energies is 0.96, 0.97, and 0.71 for the FI, MF, and O(∞) models, respectively. B) Distributions of Pearson correlation coefficients between inferred and true interaction energies across 50 random systems (each consisting of n=50 enhancers with seven binding sites) for *σ* = 0.5, 1, 1.5.

### 3.3 Finding interactions for an enhancer measured in different conditions

In the previous sections, we considered enhancers in a single cellular condition with a fixed set of known binding sites. Equivalently, instead of varying the enhancer sequences, one could imagine having a single enhancer sequence, but measured in a variety of cellular conditions (i.e. *µ*_*i*_ changes). The inference methods should apply equally well to such data for the determination of the interaction energies between TFs in the given enhancer. Developmental systems provide a natural setting for such data, where the output of a given enhancer can be measured across different cellular contexts with varying TF concentrations (i.e. chemical potentials). Before applying our inference method to such experimental data in the next section, we first test it on simulated data in this section.

We assume that the Pol-II binding probability is proportional to the measured fluorescence intensity of the target gene product. In addition, the fluorescence intensities of TFs (related to their chemical potentials, see Sec. 2.6) have also been measured, for example by imaging the fluorescence of each TF in each cell. The binding energies of TFs (*h*_*i*_) are free parameters to be inferred. We assume that the concentration of Pol II does not change across conditions; therefore, its chemical potential is set to zero.

To test our models’ ability to infer interactions from this type of data, we simulated data using the fully interacting model for an enhancer containing seven binding sites for known TFs and Pol-II. The interaction energies *J*_*i j*_, chemical potentials, and binding energies were drawn from a normal distribution with zero mean and standard deviation *σ* . For every simulated enhancer, the binding energy of the promoter was set to zero as an energy reference. For a given enhancer, we generated *M* = 1000 different cellular conditions and simulated a single output corresponding to each condition (see Sec. 2.4).

We applied all three inference methods to the simulated data. Fig. 5A shows an example of the fit, comparing the inferred *J* to the true interaction matrix, and the true binding energies to the inferred ones (the last row of the heatmaps), where *σ* = 1. The Pearson correlation coefficient in this example is 0.94, 0.85, and 0.73 for the FI, MF, and O(∞) models, respectively.

**Figure 5:**
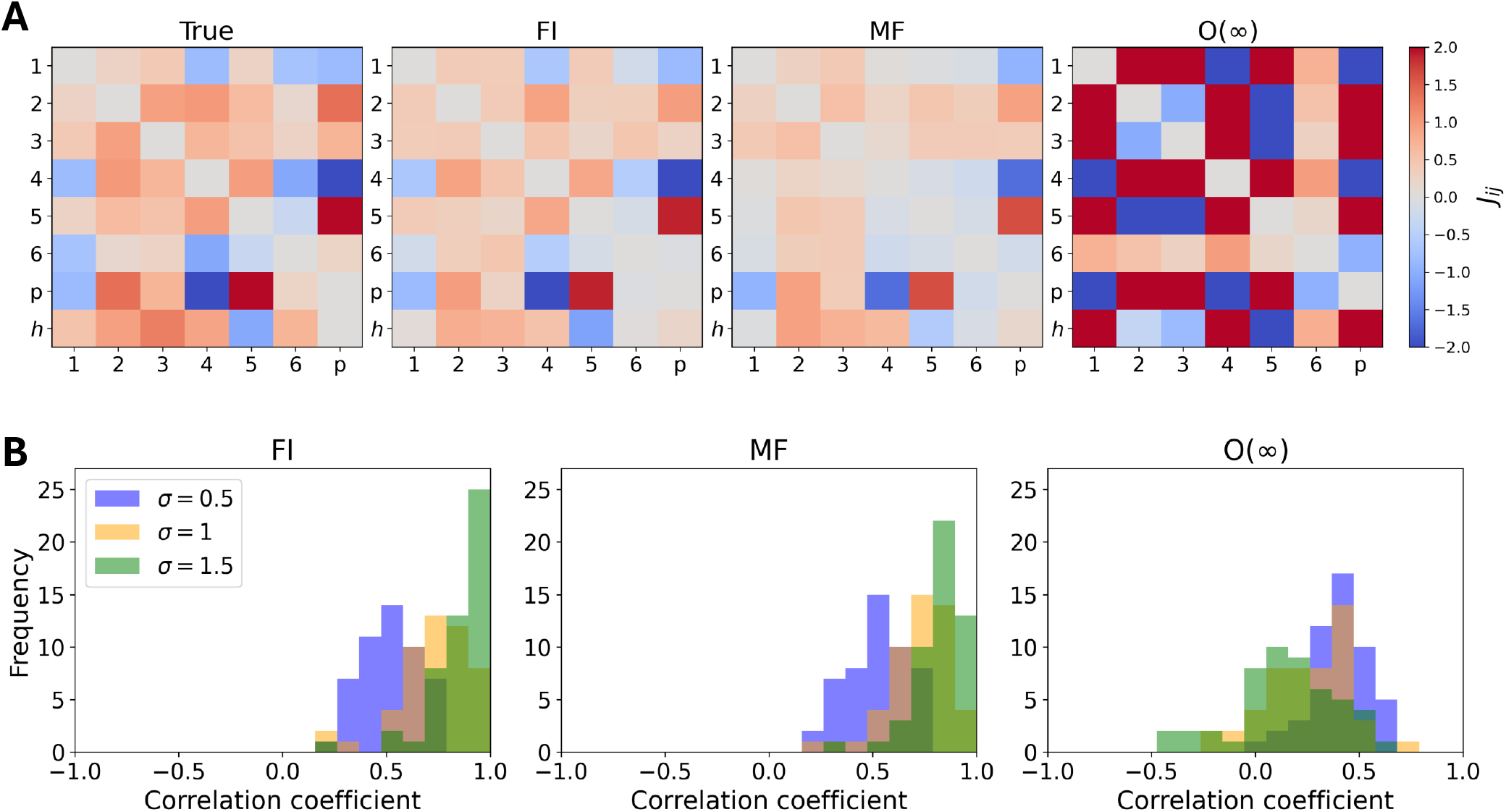
Inference of interaction energies from simulated expression data from a single enhancer measured in *M* = 1000 different cellular conditions. A) Heatmaps of the true interaction energy matrix alongside the inferred interaction and binding energies using the FI, MF, and O(∞) models, where *σ* = 1. The Pearson correlation between the inferred and true energies is 0.94, 0.85, and 0.73 for the FI, MF, and O(∞) models, respectively. B) Distributions of Pearson correlation coefficients between inferred and true interaction and binding energies across 50 random systems (each consisting of a single enhancer measured in *M* = 1000 different cellular conditions) for *σ* = 0.5, 1, 1.5.

We repeated this procedure for 50 independent random systems generated using different values of *σ* for the interaction matrices, binding energies, and chemical potentials. Consistent with the results of the previous section, the accuracy of the inference decreases as *σ* increases for the O(∞) model, as can be seen in Fig. 5B. However, for the other two inference methods the accuracy increases with *σ* . We also generated background distributions using 10000 random matrices with the corresponding *σ* values (see Fig. S3C).

### 3.4 Application of the inference method to the eve stripe-2 enhancer

One of the canonical developmental enhancers is the eve stripe-2 enhancer (eve2) from D. melanogaster. It has binding sites for the gap genes: Hb, Gt, Kr, and the maternal morphogen Bicoid (Bcd). Although well studied, and while the overall function of these proteins on eve2 expression is known (e.g. Hb, Bcd are activators and Gt, Kr are repressors [38, 39]), the significance of cooperative or inhibiting interactions between the TFs is still a matter of open investigation [34, 40–43].

In principle, these TFs bind to many different sites within the eve2 enhancer, forming a large and complex gene regulatory network. Here, we approximate this complex system using a simplified enhancer model containing only five effective binding sites (Hb, Gt, Kr, Bcd, and Pol-II), each represents the collective contribution of its corresponding TF binding sites in the original system. When fitted to experimental data, the inferred interaction and binding energies in this reduced model can be interpreted as effective energies that summarize the behavior of the full regulatory system.

As discussed in Sec. 2.3.2, the available experimental data provide only fluorescence intensities of the gap gene proteins, which are related to their chemical potentials (see Sec. 2.6). Since the number of TFs considered here is not too large, we employed the FI model, as our most accurate model, to infer effective interactions in eve2.

Before fitting the data, we excluded the TF Knirps (Kni). Although it is generally important in the regulation of the eve gene, it does not play a major role in eve2 expression [39, 44]. Moreover, its concentration is nearly constant at the location of the second stripe along the embryo (see Fig. 1D), where the model is fitted. We estimated the eve Pol-II binding probability at position *x* in the second stripe by normalizing the fluorescence intensity to the maximum eve2 intensity. We then computed the chemical potential for each factor (Fig. 6B), using as reference the intensity of each factor at the position indicated by the black star in Fig. 6B. The reference was chosen such that it yields a value closest to *m*_*p*_ = 0.5 (see Sec. 2.6).

**Figure 6:**
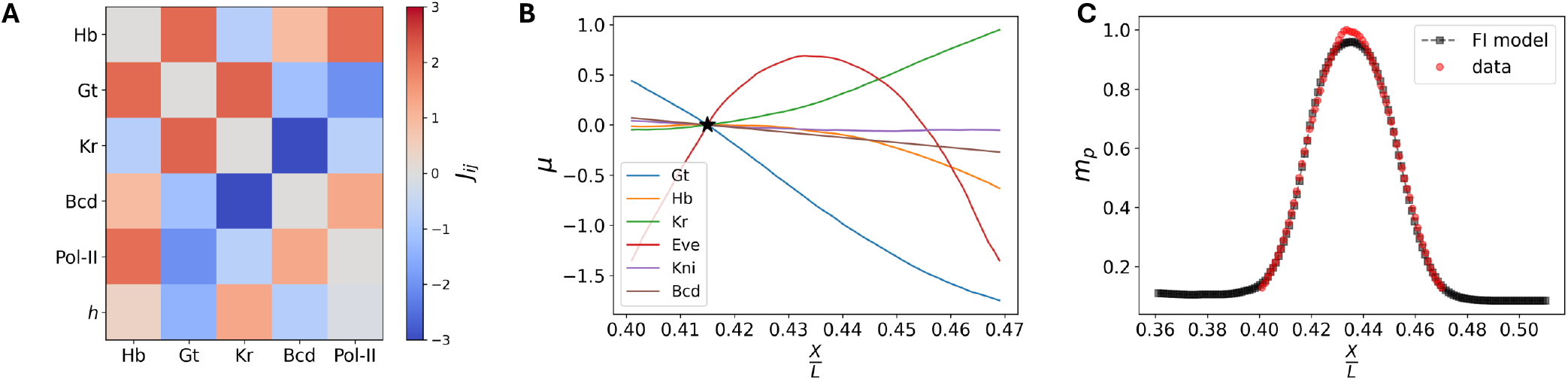
Fit results from applying the FI model to the eve2 enhancer data. A) Heatmap of the inferred interaction and binding energies. B) The chemical potentials of the TFs, used as inputs to the model at the posiion of the second stripe. C) Model fit to the estimated eve2 Pol-II binding probability derived from the data.

The inferred interaction map (Fig. 6A) shows the interaction energies among the four TFs and Pol-II at the eve2 enhancer. These interactions were obtained by fitting the FI model to the chemical potentials of the TFs (Fig. 6B) and the corresponding estimated average occupancy of Pol-II (*m*_*p*_, Fig. 6C), where regularization was employed to avoid overfitting (see S.M. 1.4). Using the inferred parameters and the chemical potentials in the flanking regions, we extrapolated *m*_*p*_ outside the stripe. The low extrapolated values of *m*_*p*_ in the flanks suggest that eve expression outside the second stripe should remain limited, consistent with the expected silencing of the eve2 enhancer beyond the second stripe.

As expected from previous studies, the inferred interaction energies between Hb, Bcd, and Pol-II are positive in eve2, indicating their activating roles, whereas the interaction energies associated with Kr and Gt with Pol-II are negative, reflecting their repressive functions. Beyond recovering these known interactions, the inferred map also reveals cooperative effects between the TF pairs Hb–Bcd, Hb–Gt, and Gt–Kr, as their inferred interaction energies are positive. In contrast, the interactions between Hb–Kr, Gt–Bcd, and Kr–Bcd are negative, indicating antagonistic relationships among these pairs. Furthermore, the inferred effective binding energies (the last row of the heatmap, Fig. 6A) suggest that Hb and Kr have stronger effective binding sites than Gt and Bcd.

## 4 Discussion

In this work, we developed a framework to infer TF–TF and TF–Pol-II interactions from gene expression data, including assays such as STARR-seq (mRNA read counts for fragments spanning an enhancer) and fluorescence-based measurements (the same enhancer across different cellular conditions). We use a biophysical Ising-type model of transcription, in which regulatory elements are characterized by site-specific binding energies (*h*_*i*_), pairwise interaction energies (*J*_*i j*_), and chemical potentials that reflect TF concentrations under different cellular conditions. Within this framework, the model computes the Pol-II binding probability, which serves as a proxy for expression.

Because the fully interacting model becomes computationally intractable for complex enhancers, we developed a hierarchy of approximations based on the mean-field representation of the model that progressively reduce computational complexity. We evaluated these approximations on simple simulated enhancer architectures and found that the FI, MF, and O(∞) models recover the ground-truth interactions used to generate the data by maximizing the likelihood for STARR-seq count data. Moreover, the first-order model (O(1)) captures effective interaction energies between TFs and Pol-II, reflecting the overall regulatory role of each factor in transcription.

We then focused on the fully interacting (FI), mean-field (MF), and all-order (O(∞)) models and applied them to simulated STARR-seq data for enhancers with randomly assigned interactions. We observed that the inferred interactions are correlated with the true interactions across all three models. However, the accuracy of the O(∞) model decreases as the magnitude of the true interaction strengths increases (i.e., as *σ* increases), whereas the accuracy of the FI and MF models does not change considerably. This behavior is in line with the assumptions underlying the approximation, which is most accurate in the weak-interaction regime. Furthermore, we observed that the inference of *J* is more accurate when the data consist of measurements across multiple enhancers in the same cellular condition, compared to a single enhancer, even when the total number of fragments is the same across the two settings.

We next simulated the case of a single enhancer across varying cellular conditions, where TF chemical potentials change, analogous to the eve2 fluorescence dataset. The method successfully inferred the interaction map between regulatory elements by minimizing the prediction error. However, the inference quality of the O(∞) model decreases as the magnitude of the changes in chemical potentials and interactions between TFs increases (i.e., with increasing *σ*). This decline in performance is expected, as the approximation relies on a Taylor expansion around a reference point, and its accuracy decreases as the system deviates further from that point. Surprisingly, however, the performance of the FI and MF models improves with increasing *σ* .

After validating the framework on simulated data, we applied the FI model to experimental fluorescence data for the eve stripe-2 enhancer. The inferred interactions agree with well-established biological knowledge [38, 39]: Hunchback and Bicoid act as activators with positive interactions with Pol-II, while Giant and Krüppel exhibit repressive roles. Beyond these known effects, our results suggest repressive interactions between the pairs Hb–Kr, Gt–Bcd, and Kr–Bcd, as well as cooperative activating interactions between Hb–Bcd, Hb–Gt, and Gt–Kr. To our knowledge, such interaction patterns are less well characterized. Moreover, the inferred binding energies suggest that Kr has the strongest effective binding strength, followed by Hb, Pol-II, Bcd, and Gt in decreasing order.

It is known that the gap genes and the morphogen Bicoid have multiple binding sites within the eve2 enhancer. In this work, however, we considered a single effective site per factor, representing the aggregate contribution of all binding sites for that TF. Consequently, the inferred interaction map reflects the collective regulatory role of each factor at the enhancer level. An extension would be to model all known binding sites explicitly and apply the framework at higher resolution. Furthermore, the model does not account for self-regulation or self-cooperativity among TFs, which can introduce non-linearities in the regulatory response that might not be captured by the model. As a result, some of the inferred interaction energies may reflect these unmodeled effects rather than true pairwise TF-TF interactions.

