## Supporting information for "Inference of enhancer-specific transcription factor interactions from gene expression data using a biophysical model"

### 1 Supporting Material

#### 1.1 Summary of parameters used in simulations

For all simple enhancer architectures, $r_0 = 1000$.

- Helper-activator

$$\vec{h} = \begin{bmatrix} -1.5 \\ -1.5 \\ -1.5 \end{bmatrix} \quad J = \begin{bmatrix} 0 & +3 & 0 \\ +3 & 0 & +3 \\ 0 & +3 & 0 \end{bmatrix} \quad \vec{\mu} = \begin{bmatrix} 0.5 \\ 0.5 \\ 0.5 \end{bmatrix}$$

- Inhibitor-Repressor

$$\vec{h} = \begin{bmatrix} 0.5 \\ 0.5 \\ 0.75 \end{bmatrix} \quad J = \begin{bmatrix} 0. & -2 & 0. \\ -2 & 0. & -2 \\ 0. & -2 & 0. \end{bmatrix} \quad \vec{\mu} = \begin{bmatrix} -0.25 \\ -0.25 \\ -0.25 \end{bmatrix}$$

- Dual-Repressor

$$\vec{h} = \begin{bmatrix} 1 \\ 1 \\ 1.5 \end{bmatrix} \quad J = \begin{bmatrix} 0. & 2 & -3 \\ 2 & 0. & -3 \\ -3 & -3 & 0. \end{bmatrix} \quad \vec{\mu} = \begin{bmatrix} -0.5 \\ -0.5 \\ -0.5 \end{bmatrix}$$

For simulations of a single enhancer tiled with $M = 1000$ fragments (Fig. 3) in Sec. 3.2, $h_i + \mu_i$ was drawn from $\mathcal{N}(0,\sigma)$ and assumed to be known.

For simulations of multiple enhancers in the same condition (Fig. 4) in Sec. 3.2, $h_i$ for each enhancer and $\mu_i$ were separately drawn from $\mathcal{N}(0,\sigma)$. $h_i$ was assumed to be known, but $\mu_i$ was treated as a free parameter and inferred by the model. $h_p$ and the chemical potential of Pol II were set to zero.

For simulations of a single enhancer measured in $M = 1000$ different cellular conditions (Fig. 5) in Sec. 3.3, $\mu_i$ and $h_i$ were each drawn from $\mathcal{N}(0,\sigma)$. $\mu_i$ was assumed to be known, but $h_i$ was treated as a free parameter and inferred by the model. $h_p$ and the chemical potential of Pol II were set to zero.

In all of the above simulations, $r_0 = 1000$.

#### 1.2 Linear Regression

Eq. 15 in the main text can be reformulated using a different set of $N$ binary features, $X_i^\nu$, defined over the entire enhancer. Specifically, we define $X_i^\nu = 1$ if site $i$ is present in fragment $\nu$, and $X_i^\nu = 0$ otherwise. With this definition, Eq. 15 can be rewritten in terms of these features, where the summation now runs over all sites $N$ in the original enhancer,

$$\left(m_p^{\nu,\alpha}\right)_{\mathrm{O}(1)} = m_{0,p}^{\alpha} + \frac{1}{4}\sum_{i=1}^{N} J_{ip} m_{0,i}^{\alpha} X_i^{\nu}, \tag{1}$$

which is a linear model. Eq. 1 can be written in the linear regression form

$$\left(m_p^{\nu,\alpha}\right)_{\mathrm{O}(1)} = b + \sum_{i=1}^{M} W_i X_i^{\nu}, \tag{2}$$

where

$$W_i = \frac{1}{4} J_{ip} m_{0,i}^{\nu,\alpha}, \qquad b = m_{0,p}^{\alpha}. \tag{3}$$

More compactly, for the entire dataset, this equation can be written in vector form as

$$\left(\vec{m}_p^{\alpha}\right)_{\mathrm{O}(1)} = b\vec{1} + X\vec{W}, \tag{4}$$

where $X$ is an $M \times N$ binary feature matrix, $\left(\vec{m}_p^{\alpha}\right)_{\mathrm{O}(1)}$ is the vector of Pol-II occupancies for all fragments, and $\vec{1}$ is an $M$-dimensional vector of ones. As explained in the main text, the vector $\vec{W}$ is obtained via a Poisson regression fit. Once $W_i$ is determined, the effective interaction energy between TF $i$ and Pol-II is given by

$$J_{ip} = \frac{4W_i}{m_{0,i}^{\alpha}}. \tag{5}$$

### 1.3 Background distributions

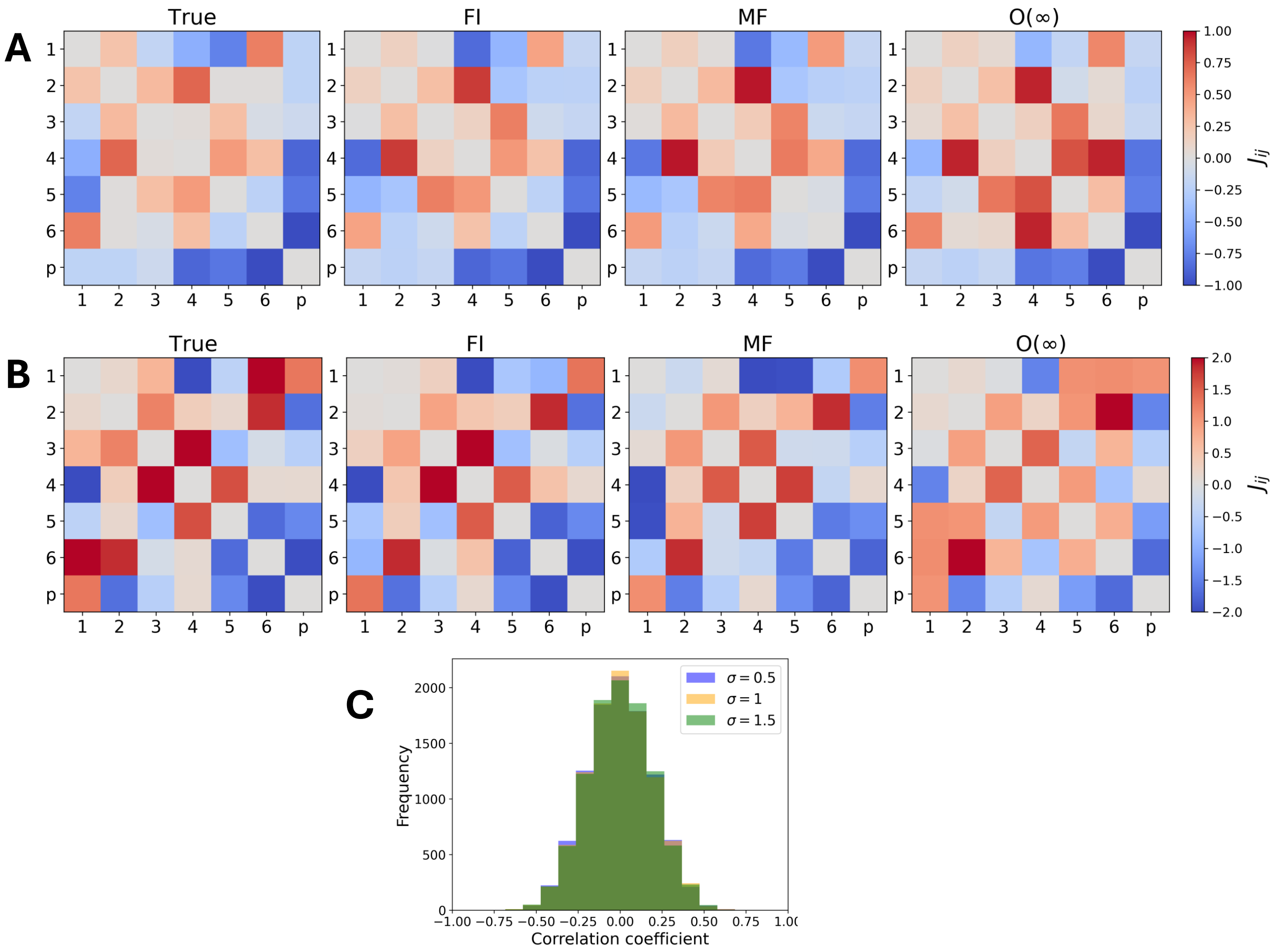


Figure S1: Inference of interaction energies for random complex enhancers using $M = 1000$ random fragments each. A) Heatmaps of the true interaction energy matrix alongside the inferred interaction energies using the FI, MF, and O($\infty$) models for a system with seven binding sites, where $\sigma = 0.5$ and $\sigma = 1.5$ in A and B, respectively. The Pearson correlation coefficients between the inferred and true energies are 0.94, 0.93, and 0.90 for FI, MF, and O($\infty$) models, respectively in panel A, and 0.87, 0.85, and 0.77 in panel B. C) Background distribution of correlation coefficients between the true matrices shown in A, B, and Fig. 3A, where each correlation coefficient was computed between the true matrix and 10000 randomly generated matrices with the same $\sigma$.

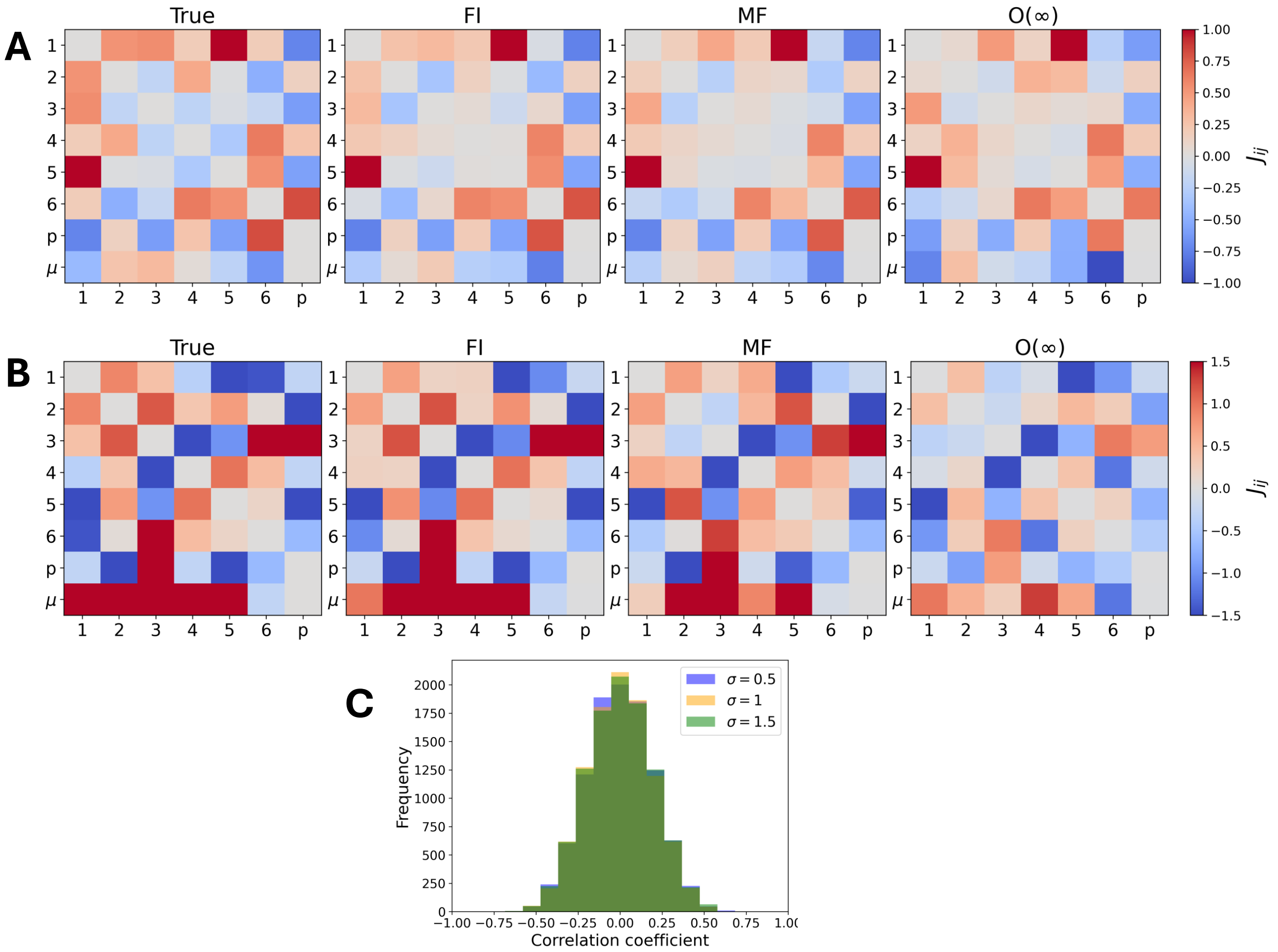


Figure S2: Inference of interaction energies from systems with multiple enhancers (n=50), where each enhancer has varying-strength binding sites for the same TFs, and is randomly tiled with 20 fragments. Heatmaps of the true interaction energy matrix alongside the inferred interaction energies and chemical potentials using the FI, MF, and O($\infty$) models, where $\sigma = 0.5$ and $\sigma = 1.5$ in A and B, respectively. The Pearson correlation coefficients between the inferred and true energies are 0.94, 0.93, and 0.88 for FI, MF, and O($\infty$) models, respectively in panel A, and 0.99, 0.93, and 0.83 in panel B. C) Background distribution of correlation coefficients between the true matrices shown in A, B, and Fig. 4A, where each correlation coefficient was computed between the true matrix and 10000 randomly generated matrices with the same $\sigma$.

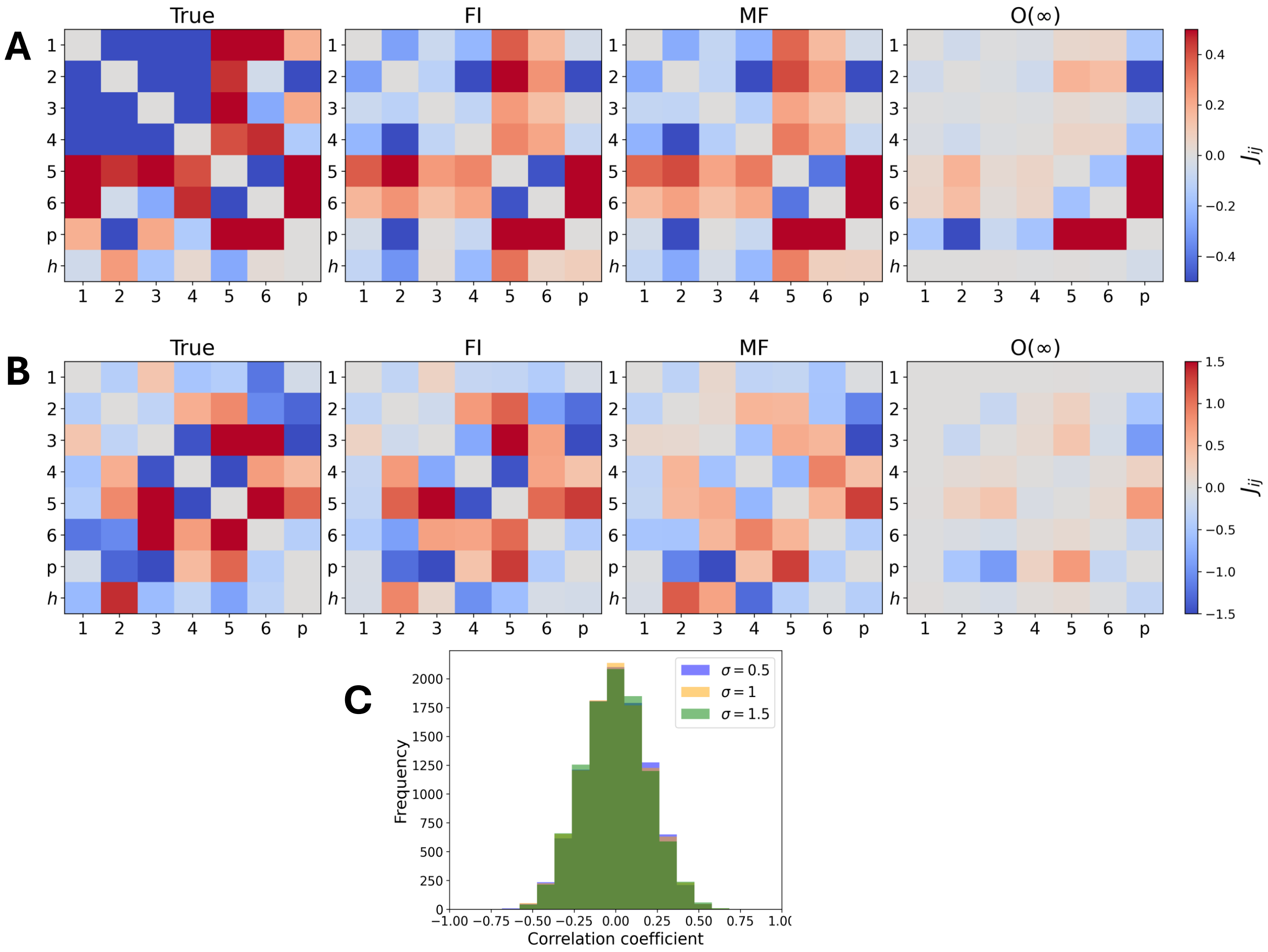


Figure S3: Inference of interaction energies from simulated expression data from a single enhancer measured in $M = 1000$ different cellular conditions. A) Heatmaps of the true interaction energy matrix alongside the inferred interaction and binding energies using the FI, MF, and O(∞) models, where $\sigma = 0.5$ and $\sigma = 1.5$ in A and B, respectively. The Pearson correlation coefficients between the inferred and true energies are 0.74, 0.75, and 0.60 for FI, MF, and O(∞) models, respectively in panel A, and 0.92, 0.76, and 0.57 in panel B. C) Background distribution of correlation coefficients between the true matrices shown in A, B, and Fig. 4A, where each correlation coefficient was computed between the true matrix and 10000 randomly generated matrices with the same $\sigma$.

#### 1.4 Regularization

For the eve2 enhancer data, our FI model is prone to overfitting, particularly due to the limited number of data points (70 in total) for training. To mitigate this, we added an $\ell_2$ regularization term to the loss function. To determine the regularization constant, $\lambda_r$, we performed 5-fold cross-validation, with the data randomly shuffled before splitting into folds.

The resulting validation loss is shown in Fig. S4. The selected value of the regularization parameter ($\lambda_r^* = 0.1$) is indicated by the red star.

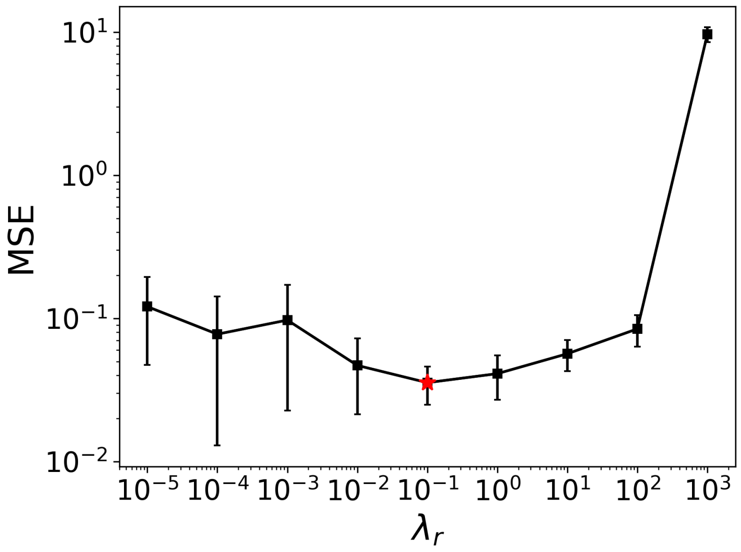


Figure S4
